# Pioneer activity of an oncogenic fusion transcription factor at inaccessible chromatin

**DOI:** 10.1101/2020.12.11.420232

**Authors:** Benjamin Sunkel, Meng Wang, Stephanie LaHaye, Benjamin J. Kelly, James R. Fitch, Frederic G. Barr, Peter White, Benjamin Z. Stanton

## Abstract

Recent characterizations of pioneer transcription factors have led to new insights into their structures and patterns of chromatin recognition that are instructive for understanding their role in cell fate commitment and transformation. Intersecting with these basic science concepts, the identification of pioneer factors (PFs) fused together as driver translocations in childhood cancers raises questions of whether these fusions retain the fundamental ability to invade repressed chromatin, consistent with their monomeric PF constituents. In this study, we define the cellular and chromatin localization of the translocation, PAX3-FOXO1, an oncogenic driver of childhood rhabdomyosarcoma (RMS), derived from a genetic fusion of PFs. To quantitatively define its chromatin-targeting functions and capacity to drive epigenetic reprogramming, we developed a new method for ChIP-seq with per-cell normalization (pc-ChIP-seq). Our quantitative localization studies address structural variation in RMS genomes and reveal novel insights into heterochromatin localization of PAX3-FOXO1. From these studies, we report novel pioneer function for the major driver oncogene in RMS, with repressed chromatin binding and nucleosome-motif targeting in human cells.

## Introduction

The pioneer factor subset of the broad transcription factor (TF) class of proteins, possesses conserved motifs common to TFs, including a structured DNA-binding domain (DBD) responsible for motif recognition and a flexible transactivation domain mediating regulated recruitment (Ptashne and Gann, 1997). Distinct structural elements of pioneer factors (PFs) provide a unique capacity for high-affinity DNA binding despite steric deterrents at the chromatin interface, including nucleosomes. However, the combinatorial rules or “logic” of domain organization within PFs, requisite to achieve nucleosomal-motif recognition, remain obscure (Fernandez Garcia et al., 2019). Moreover, it is presently unknown if two pioneer factors fused together will possess pioneer activity in the resulting chimera.

Among characterized PFs, the forkhead family has been studied extensively at the structural and regulatory levels in mammalian development and also in multiple cancers (Herman et al., 2021). A winged-helix DNA-binding domain is conserved across individual members of this protein family. In the case of FOXA1, the winged-helix domain mimics the structural features of the linker histone H1, disrupting H1-compacted chromatin together with the FOXA1 C-terminal domain (Cirillo et al., 2002; Cirillo et al., 1998; Zhou et al., 2020). Consistent with its high degree of sequence and structural similarity, the FOXO1 protein also recognizes its cognate DNA sequence motif within H1-compacted nucleosome arrays, initiating local DNase hypersensitivity through disruption of histone:DNA contacts without input from chromatin remodelers (Hatta and Cirillo, 2007). Importantly, chromatin decompaction following nucleosomal motif recognition by PFs is not necessarily associated with nucleosome eviction (Cirillo et al., 2002; Hatta and Cirillo, 2007). In a recently reported example, FOXA2 binding was shown principally to mediate the induction of nucleosome spacing within tissue-specific *cis*-regulatory regions (Iwafuchi-Doi et al., 2016). Findings such as these reinforce that PFs are capable of decompacting and stably binding to nucleosome-occupied regions of the genome, while recruitment of additional factors may be necessary for the formation of active and accessible regulatory elements generally associated with gene activation.

While evidence for direct nucleosomal motif recognition by putative PFs continues to emerge (Fernandez Garcia et al., 2019; Zhu et al., 2018), additional compact chromatin binding behaviors of PFs such as heterochromatin recognition and mitotic chromatin bookmarking are emerging from cell imaging studies. In addition to forkhead factors FOXI1 and FOXA1 (Yan et al., 2006; Zaret et al., 2008), other pioneer factors including SOX2, OCT4, and PAX3 are retained on compact mitotic chromatin (Deluz et al., 2016; Teves et al., 2016; Wu et al., 2015). In the case of SOX2, this may serve to mark specific genes for post-mitotic reactivation, while mitotic chromatin binding by PAX3 may instead be related to its reported function in the stable repression of microsatellite transcription via establishment and maintenance of H3K9me3-marked heterochromatin domains across cell divisions (Bulut-Karslioglu et al., 2012). These fundamental molecular functions of PFs likely underlie their central role in *de novo* activation of lineage-defining gene expression programs during tissue differentiation and contribute to heritable transmission of these gene programs during development.

In alveolar rhabdomyosarcoma, two pioneer factors, PAX3 and FOXO1, are fused in-frame in the recurrent translocation between chromosome arms 2p and 13q (Galili et al., 1993). The resulting PAX3-FOXO1 fusion is an oncogenic driver that has been described as binding active regulatory elements alongside myogenic TFs (Gryder et al., 2019), while its nucleosome targeting function in inactive or repressed chromatin domains remains unstudied. Neither retention of canonical pioneer activity nor the emergence of functions distinct from the wild-type PAX3 or FOXO1 monomers has been rigorously defined for PAX3-FOXO1 in fusion-positive rhabdomyosarcoma (FP-RMS). Given the relatively low mutational frequencies in FP-RMS, which can be approximated at 0.1 protein-coding mutations per Mb (Shern et al., 2014), we hypothesized that the pioneer function of PAX3-FOXO1, defined by targeting to nucleosomal motifs within inaccessible chromatin, might underlie its transforming potential in this tumor. However, the mechanisms through which PAX3-FOXO1 engages distinct classes of chromatin have remained poorly understood.

The cascade of initiation events for a tumor are unlikely to involve mere stabilization of pre-existing transcriptional networks or DNA-accessibility, but rather, a restructuring of regulatory elements into a new state that differs from a cell of origin (Stanton et al., 2017; Suva et al., 2013). Presently, the cell of origin for FP-RMS remains unknown. Expression of PAX3-FOXO1, along with other highly expressed TFs in FP-RMS, is reminiscent of both neuronal tissue and developing muscle (Galili et al., 1993; Raghavan et al., 2019; Yu et al., 2004), contributing to ambiguity in defining a tissue of origin. Often, transcriptional reprogramming represents the functional output of tissue-specific pioneer factors (Takahashi and Yamanaka, 2006; Vierbuchen et al., 2010). It is noteworthy that tumors prevalent in children are frequently defined by a profound failure of cellular differentiation (Nacev et al., 2020). In these and other relatively low-mutation-burden tumors, there has been increasing evidence suggesting that disruption of transcription factors drives reprogramming into altered epigenetic states. The role of PFs like SOX2, PAX3, and FOXO1 in developmental reprogramming may be analogous to pioneer activity in establishing cell-fate decisions in pediatric cancers, including synovial sarcoma and MPNST (Kadoch and Crabtree, 2013; Miller et al., 2009), where mis-regulation of SOX-family pioneer factors occurs, as well as in FP-RMS, where PAX3 and FOXO1 are frequently fused as a chimeric oncoprotein.

In the present study, we address the possible retention of PF activity in the PAX3-FOXO1 fusion oncoprotein at the chromatin level, as a basis for understanding its role in FP-RMS initiation. Well-established features of *bona fide* PFs guide our investigation: 1) binding to repressed/compact/inaccessible chromatin, 2) nucleosomal motif recognition and occupancy. Our biochemical and high-resolution genomic analyses reveal steady-state association of PAX3-FOXO1 with heterochromatin features, while kinetic studies of PAX3-FOXO1 induction reveal rapid targeting of PAX3-FOXO1 to nucleosome occupied regions where the fusion is retained often without inducing accessibility. These findings reveal an interplay between PAX3-FOXO1 and H3K9me3 domain patterning, opening new avenues for further understanding the chromatin level role of PAX3-FOXO1 in heritable transmission of oncogenic events in FP-RMS.

## Results and Discussion

### Cellular and genomic localization of the PAX3-FOXO1 fusion transcription factor to inactive chromatin

We asked whether the PAX3-FOXO1 fusion in rhabdomyosarcoma exhibited fundamental properties of known pioneers. Homology analysis of PAX3 and FOXO1 amino acid sequences demonstrated that evolutionarily conserved residues within the PAX3 paired domain and homeobox domain are completely retained in the fusion TF (**Figure 1A**). While the conserved nuclear-export and -localization sequences (NES and NLS), as well as the transactivation domain (TAD) of FOXO1 remain intact in the fusion, the forkhead or winged-helix domain is N-terminally truncated. The PAX3-FOXO1 fusion product retains the essential amino acid residues within alpha-helix 3 for recognition of FOXO1 DNA sequence motifs (5’-TGTTTAC-3’) and the flexible wings W1 and W2, which have been shown to stabilize FOXO1:DNA interactions through direct phosphate-backbone contacts (**Figure 1A**) (Brent et al., 2008). Despite the truncation, it was predicted by analysis of the primary amino acid sequence that the ordered DNA-binding domains and the disordered transactivation domains, originating from both PAX3 and FOXO1, are retained in the resulting fusion product (**Figure S1A**). As the PAX3-FOXO1 chimera retains many of the conserved features of pioneer protein families, we went on to systematically assess the ability of this fusion TF to initiate reprogramming at the chromatin level to better understand epigenetic initiation in FP-RMS.

**Figure 1.**
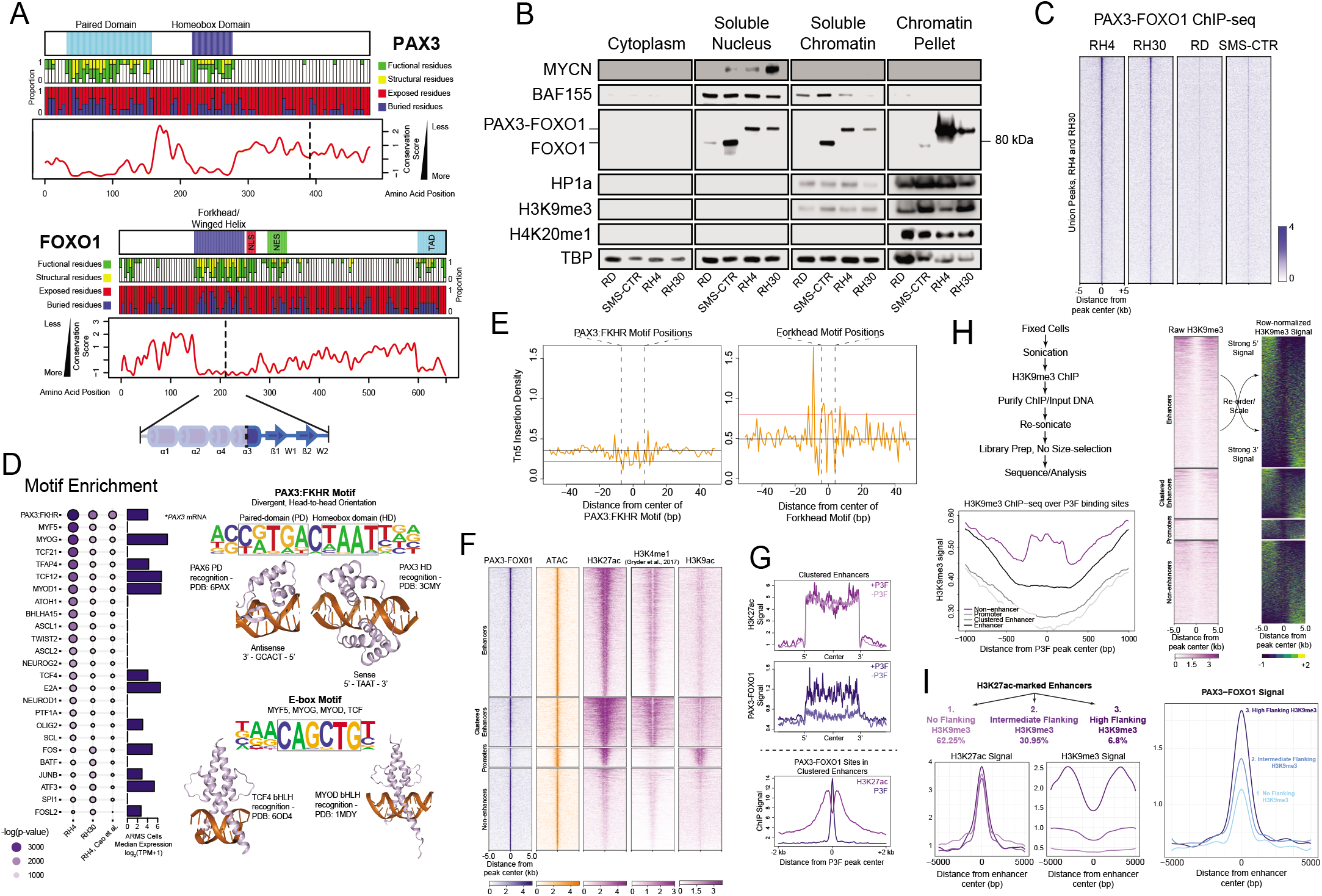
Cellular and genomic localization of the PAX3-FOXO1 fusion transcription factor to inactive chromatin. **(A)** Evolutionary conservation of PAX3 (top) and FOXO1 (bottom) amino acid sequences. Vertical dashed lines indicate the breakpoint position resulting in the formation of the PAX3-FOXO1 fusion. FOXO1 inset details N-terminal truncation of the winged helix domain. (**B**) FN-RMS (RD, SMS-CTR) and FP-RMS (RH4, RH30) cells were fractionated to interrogate protein localization within the cytoplasm, soluble nucleus, soluble chromatin (sonication sensitive), and chromatin pellet (sonication insensitive) compartments of the cell. (**C**) A C-terminal FOXO1 antibody was utilized to perform per-cell ChIP-seq (pc-ChIP-seq) of PAX3-FOXO1 in a panel of RMS cells. Spike-in normalized signal intensity from each cell line was displayed across the union of PAX3-FOXO1 sites identified in RH4 and RH30 cells. (**D**) HOMER motif analysis was performed on PAX3-FOXO1 binding sites from anti-FOXO1 pc-ChIP-seq (RH4 and RH30), and pFM2 ChIP-seq (RH4, Cao *et al., Cancer Res* 2010). RNA-seq expression for the transcription factor corresponding to each motif is displayed as the median expression for PAX3-FOXO1+ alveolar RMS (ARMS) cells in the Cancer Cell Line Encyclopedia (CCLE). The PAX3-FKHR motif consists of a Paired Domain (PD) and a Homeobox Domain (HD) motif arranged in a divergent, head-to-head orientation. Representative PDB structures demonstrate the recognition of PD and HD motifs by short alpha-helices, where E-box motifs are recognized by the extended alpha-helices of bHLH family transcription factors. (**E**) ATAC-seq was performed in RH4 cells, and Tn5 insertion positions were calculated by the NucleoATAC pipeline. Insertion rates are displayed with respect to high confidence PAX3-FOXO1 (left) and FOXO1 (right) motif positions (vertical dashed lines) within PAX3-FOXO1 binding sites identified using FIMO. The median insertion rate outside and within the motif positions is displayed as a black and red horizontal line, respectively. (**F**) Heatmap of PAX3-FOXO1, histone marks and ATAC-seq in P3F binding sites categorized as promoters, enhancers, clustered enhancers, and non-enhancers in RH4 cells (**G**) Analysis of H3K27ac and P3F in enhancer clusters. (Top, Middle) H3K27ac (Top) and P3F (Middle) signal over clustered enhancer regions with or without an overlapping P3F binding site. (Bottom) P3F and H3K27ac signal centered over P3F binding sites occurring within enhancer clusters. (**H**) H3K9me3 ChIP-seq was performed in RH4 cells with a modified library preparation. H3K9me3 profiles were characterized over P3F binding sites in promoters, enhancers, clustered enhancers, and non-enhancers. Ordering P3F sites according to the degree of 5’-3’ H3K9me3 signal imbalance, H3K9me3 signals were found to exhibit asymmetry around P3F binding sites. (**I**) H3K27ac-marked enhancer regions were divided into three categories displaying 1) no flanking H3K9me3, 2) intermediate flanking H3K9me3, or 3) high flanking H3K9me3. H3K27ac, H3K9me3, and P3F profile plots were generated for each enhancer category. Up- and down-stream H3K9me3 imbalance were significantly higher in “high flanking H3K9me3” sites than in “intermediate flanking H3K9me3” sites (Up-stream p-value = 1×10^-7^, Down-stream p-value = 0) and “no flanking H3K9me3” sites (Up-stream p = 0, Downstream p-value = 0) (Anova, Tukey’s HSD correction).

We were first motivated to address the enigmatic question of whether the PAX3-FOXO1 driver oncoprotein can initiate chromatin reprogramming, as it has only been previously observed to bind active, accessible chromatin. Thus, we tested whether PAX3-FOXO1 genomic localization is distinct from non-pioneer TFs and whether it has the capacity to bind regions with lower levels of accessibility. We conducted a genomewide correlation between various chromatin factors and PAX3-FOXO1 using ChIP-seq datasets from the FP-RMS cell line RH4, available publicly or generated for this study (Cao et al., 2010; Gryder et al., 2017). Our results revealed remarkable dissimilarity between PAX3-FOXO1 binding and localization of FP-RMS <u>c</u>ore regulatory <u>TF</u>s (CRTFs; MYCN, MYOG, MYOD1), chromatin structural components (CTCF, RAD21), enhancer binding/regulatory factors (MED1, BRD4, p300), active histone modifications (H3K27ac, H3K9ac, H3K4me1), accessibility (ATAC-seq), and SWI/SNF chromatin remodeling complex subunits (BRD9, DPF2) (**Figure S1B**). Remarkably, genome-wide PAX3-FOXO1 signal showed the second strongest correlation with the heterochromatic histone modification, H3K9me3, behind the repressive H3K27me3 modification. This observation revealed a unique genomic binding preference for PAX3-FOXO1 compared to other CRTFs in FP-RMS cells and suggested PAX3-FOXO1 may substantially reside in inactive regions of the genome.

To determine whether the pattern of PAX3-FOXO1 occupancy we observed could be attributed to its localization outside the active, euchromatic nuclear compartment, we employed a stringent, sequential fractionation protocol. We aimed at distinguishing cellular components associated with increasingly insoluble compartments of the nucleus in a panel of RMS cells, including PAX3-FOXO1 fusion-negative cell lines (RD, SMS-CTR) and fusion-positive cell lines (RH4, RH30). We found that the euchromatic SWI/SNF subunit BAF155 and the CRTF, MYCN, were readily extracted from the chromatin fiber in all cells when exposed to 500 mM NaCl extraction buffer (soluble nucleus; **Figure 1B**). Residual chromatin-bound BAF155 was almost exclusively observed in the sonication-sensitive chromatin fraction (soluble chromatin; **Figure 1B**). Consistent with previous characterization of this euchromatic fraction (Becker et al., 2017), we found that it was largely depleted of constitutive heterochromatin components such as HP1α as well as repression- and compaction-related histone modifications such as H3K9me3 and H4K20me1 (Becker et al., 2017; Hori et al., 2014). TATA-binding protein (TBP), with its dual role in active gene transcription and mitotic bookmarking was readily observed in all nuclear and chromatin fractions including the residual insoluble chromatin pellet (**Figure 1B**) (Teves et al., 2018). Blotting with an antibody recognizing the C-terminus of FOXO1 (epitope retained in the PAX3-FOXO1 fusion), we observed nuclear staining for wild type FOXO1 in RD and SMS-CTR cells that was largely excluded from the insoluble chromatin fraction. Unexpectedly, nuclear PAX3-FOXO1 in RH4 and RH30 cells, was readily observed in the sonication-resistant, insoluble portion of the chromatin (**Figures 1B** and **S1C**). Thus, the PAX3-FOXO1 fusion protein appears capable of invading compact, repressed chromatin. This behavior, necessary for PF-mediated cell fate transition, is an unexplored feature of this oncogenic factor, previously unknown in this tumor.

To further delineate the unique chromatin binding features of PAX3-FOXO1 in FP-RMS, we next sought to define its genome-wide occupancy profile by optimizing high-specificity, spike-in normalized ChIP-seq conditions in RH4 cells using a C-terminal FOXO1 antibody (**Figure S2A**). Our method, which we have called per-cell ChIP-seq (pc-ChIP-seq), addresses global changes in ChIP signal resulting from differential chromatin content or output between cell lines and treatment conditions, by introducing known ratios of mouse spike-in cells prior to sonication (Gryder et al., 2020). We justified this methodology by analyzing input sequencing libraries to reveal vastly different relative genome sizes across RMS cell lines (Range: 5.25-10.02 Gb, see **Methods**). Upon sequencing, we benchmarked our data against previously published PAX3-FOXO1 ChIP-seq generated with a non-commercial antibody, pFM2 (epitope spanning the fusion breakpoint) (Cao et al., 2010). We found that our ChIP-seq conditions identified 69% of reported binding events (**Figure S2B**), and with improved signal, we revealed 7,341 novel and high-strength PAX3-FOXO1 sites (**Figures S2B–S2C**). Employing identical ChIP conditions in additional RMS cell lines (**Figure S2D**), integrated analysis of our spike-in normalized anti-FOXO1 ChIP-seq revealed reproducible binding profiles that were concordant between FP-RMS cell lines (**Figure 1C**). Additionally, while fusion-negative RMS cells (FN-RMS), particularly SMS-CTR, express nuclear FOXO1 (**Figure 1B**), anti-FOXO1 ChIP-seq in these cells did not result in high confidence peak assignments (9 and 0 peaks called in RD and SMS-CTR, respectively), demonstrating that our approach is highly specific for generating PAX3-FOXO1 ChIP-seq even in the presence of FOXO1. As an internal control of FOXO1 antibody specificity in our RH4 PAX3-FOXO1 ChIP-seq dataset, we also analyzed reads aligning to the mouse genome, detecting just 135 low-confidence peaks in C2C12 cells with a greater than 14-fold reduction in the <u>F</u>raction of <u>R</u>eads <u>i</u>n Peaks (FRiP) score relative to RH4 cells. By comparing our PAX3-FOXO1 data in FP-RMS cells against nearly 26,000 publicly available ChIP-seq datasets from human tissues and cell lines using the *Cistrome DB Toolkit* (Mei et al., 2017; Zheng et al., 2019), we further supported the specificity of our approach, finding no significant correlation between PAX3-FOXO1 binding and any public wild type PAX3 or FOXO1 ChIP-seq data (**Figure S2E**). Taken together, our robust method of PAX3-FOXO1 genomic localization provides a quantitative measure of fusion protein binding across individual cell models, while suggesting distinct binding patterns for PAX3-FOXO1 relative to its wild type constituents.

We further investigated the quality and specificity of our PAX3-FOXO1 ChIP-seq by performing motif analysis. Among known motifs curated by HOMER, a “PAX3:FKHR” motif derived from previously published PAX3-FOXO1 ChIP-seq was the top enriched sequence (p-value = 10^-1625^) (**Figure 1D**) (Cao et al., 2010; Gryder et al., 2017). By convention, FKHR is synonymous with FOXO1, and we have used the “PAX3:FKHR” nomenclature throughout this study when referring to this motif from HOMER. *De novo* motif identification also returned a sequence with highest similarity to the known PAX3:FKHR motif as the most enriched (p-value = 10^-1753^). This motif is comprised of a paired-domain and a homeobox recognition sequence arranged in a divergent head-to-head orientation separated by a single cytosine. The remaining motifs enriched within PAX3-FOXO1 peaks (RH4 and RH30) were mainly E-Box motifs recognized by the non-pioneer (Fernandez Garcia et al., 2019), extended alpha-helical DBDs of basic helix-loop-helix (bHLH) family TFs such as MYF5, MYOG, MYOD1, and TCF, many of which are expressed in FP-RMS cells (**Figure 1D**) (Ghandi et al., 2019). Notably, forkhead motifs were only modestly enriched in PAX3-FOXO1 binding sites relative to background (FOXO1 motif p-value = 10^-31^), suggesting sequence-specific binding of the fusion is mediated predominantly through the intact PAX3 DBD. This was further supported by the existence of a transposase-protected footprint centered on high-confidence PAX3:FKHR motifs in RH4 binding sites, while no clear footprint was observed over forkhead motif positions by ATAC-seq (**Figure 1E**). We proceeded with high confidence in our PAX3-FOXO1 (P3F) dataset as a foundation for discovering novel genomic binding patterns and potential pioneer function of this oncogenic fusion protein.

Having identified thousands of previously unknown P3F binding sites in the FP-RMS genome, we were motivated to understand the genomic context and characteristic epigenetic state at these regions. As previously described, we found that P3F mainly occupies non-promoter, often distal intergenic regions (**Figure S2F**) (Cao et al., 2010). Linking each P3F peak to nearby transcription start sites (TSS) with GREAT (McLean et al., 2010), we found that the limited number of promoter and promoter-proximal P3F binding sites were associated with general cellular and metabolic pathways (**Figure S2G**). Interestingly, more distal P3F binding sites up to 500 kilobases from a TSS were strongly linked to genes within neurogenic differentiation processes. However, these neurogenic genes show similarly high expression across FP-RMS, FN-RMS, neuroblastoma, and glioma cell lines relative to all other models in the Cancer Cell Line Encyclopedia (CCLE) (**Figure S2H**). Thus, while our discovery of new P3F binding sites reveals associations of this fusion oncoprotein with gene pathways beyond the myogenic transcriptional circuitry, we find that P3F expression and binding alone cannot predict high expression of these genes. Further investigation of the functional consequences of P3F chromatin binding may reveal an order of events, providing clues regarding a cell-of-origin.

As in previous studies (Cao et al., 2010; Gryder et al., 2019; Gryder et al., 2017), we confirmed that P3F shows enriched occupancy of gene-regulatory enhancers compared to promoters, including 1,520 individual binding sites within 932 TSS-distal, high intensity H3K27ac clusters stitched together by ROSE (**Figures 1F** and **S3A**). However, we observed no correlation between H3K27ac strength and P3F occupancy in these sites. Clustered enhancers lacking the fusion oncoprotein exhibited nearly equal levels of H3K27ac compared to those bound by P3F (**Figure 1G**). One explanation for these observations is that P3F occupies a narrow footprint within discrete enhancer elements, and thus does not broadly influence the entire landscape of these broad regions (**Figure 1G**). However, with a general lack of correlation between P3F binding and H3K27ac enrichment across all enhancers, our results are consistent with a limited role for P3F in enhancer acetylation (**Figure S3B**). Importantly, we found that 2,565 P3F sites occur outside of enhancers and confirmed that these regions are in fact depleted of both H3K27ac and H3K4me1 while maintaining DNA accessibility (**Figure 1F**). Despite evolutionary conservation of the DNA sequences within these P3F sites (**Figure S3C**), we could not readily discern their regulatory role within the FP-RMS epigenome using publicly available datasets (**Figure S3D**). This suggests that these non-enhancer P3F peaks exhibit lineage-restricted epigenetic signatures, a feature common to promoter-distal gene-regulatory elements. To understand if these regions might be better classified as “poised,” we mapped our H3K27me3 ChIP-seq data to PAX3-FOXO1 binding sites compared to true H3K27me3-marked regions (**Figure S3E**). Given the relative lack of H3K27me3 signal, combined with depletion of both H3K27ac and H3K4me1 at these P3F-bound sites, our data does not support classifying these novel regions as enhancers on the basis of their epigenetic signatures.

We next evaluated expression changes of genes linked to each type of P3F site, including novel non-enhancer regions. Through integrative meta-analyses of our P3F-bound sites with publicly available gene expression data, we did not observe substantial changes in the expression of genes proximal to P3F binding sites associated with altered P3F expression (Ghandi et al., 2019; Gryder et al., 2017). This result was robust across conditions of P3F knockdown or add-back, as well as when comparing FP-RMS versus FN-RMS cell lines (**Figure S3F**). Subsequent analyses of RNA-seq profiles from a cohort of PAX3-FOXO1+ FP-RMS vs. FN-RMS (embryonal, ERMS, subtype) patients (Downing et al., 2012; McLeod et al., 2021), we once again found that relatively few genes associated with PAX3-FOXO1 binding were differentially expressed in patients based on PAX3-FOXO1 fusion status (**Figure S3G**). Across P3F binding site categories, less than 30% of proximal genes were differentially expressed in patient samples (average 23.7%, fold change > 2, adjusted p-value < 0.05), and these genes showed similar likelihood of being up- or down-regulated in P3F+ patients. These data indicate that P3F binding alone may be a poor predictor of gene activation in FP-RMS tumors and model systems, suggesting additional inputs are required to initiate gene expression reprogramming in a contextdependent manner.

We then asked whether the genomic occupancy profile of P3F related to our finding that the fusion oncoprotein is readily localized to the insoluble chromatin pellet in cell fractionation assays (**Figure 1B**). To this end, we performed H3K9me3 ChIP and modified our ChIP-seq library preparation protocol to achieve better sequencing coverage across relatively sonication-resistant portions of the genome (see **Methods**, **Figure 1H**) (Becker et al., 2017). Aligning our resulting data to P3F binding site categories, we found that H3K9me3 signals were specifically depleted from *enhancer, clustered enhancer,* and *promoter* binding sites, while *non-enhancer* sites were often associated with a local H3K9me3 peak directly overlapping P3F binding events (**Figure 1H**). We noted a relatively high H3K9me3 signal flanking all P3F peaks, regardless of binding site category. We assessed whether these were balanced (both up-*and* down-stream) vs. asymmetric (only up-*or* down-stream) H3K9me3 signals adjacent to P3F sites. We isolated H3K9me3 signals up-stream of P3F sites from H3K9m3 signals down-stream of P3F sites and ranked sites in each category according to their 5’-3’ imbalance. From these analyses, it was clear that the majority of P3F sites exhibit some degree of asymmetry in the adjacent H3K9me3 signal, and this occurs at each binding site category (**Figures 1H and S3H**). Importantly, we found that enhancers generally did not display a strong up- or down-stream H3K9me3 signal. In the rare occurrences that enhancers were flanked by strong, asymmetric H3K9me3 signals (6.8% of enhancers), these regions displayed relatively strong P3F occupancy (**Figure 1I)**. This analysis of H3K9me3 brings clarity to our earlier genome-wide correlation analysis, which showed only moderate correlation between P3F and H3K9me3 signals, perhaps a result of P3F most often residing adjacent to a strong H3K9me3 signal with less frequent, direct overlap. Additionally, we learn that the enrichment of P3F to the insoluble chromatin fraction of the nucleus may reflect its occupancy at the boundaries of H3K9me3 domains, corresponding to a rare subset of active regulatory elements immediately flanked by heterochromatin. Together, the results above are suggestive of a pioneer role for PAX3-FOXO1 whose binding is associated with chromatin accessibility often on the edges of repressed chromatin tracts, but which may be insufficient for chromatin activation and highly context-dependent for gene regulation.

### Early genomic targeting of PAX3-FOXO1 exhibits nucleosomal motif recognition

While binding to inactive chromatin is a feature differentiating PFs from traditional TFs, another rigorous definition of nucleosomal motif binding may be applied to understand how a TF with transforming potential is capable of invading repressed sites (Fernandez Garcia et al., 2019; Lerner et al., 2020). However, this function cannot be fully addressed at equilibrium, in which pioneer binding may in certain cases result in rapid nucleosome destabilization or eviction upon recruitment of additional factors (Yan et al., 2018). Observing this phenomenon requires kinetic regulation of PAX3-FOXO1 to monitor chromatin state changes over short time scales. To address the limitations of studying FP-RMS cells at steady state, we employed a model system of immortalized human myoblasts (Dbt), engineered with a doxycycline-inducible PAX3-FOXO1 construct (Dbt/iP3F; **Figure S4A**) (Pandey et al., 2017). With kinetic control of P3F expression, we set out to establish the immediate-early targets of P3F binding and assess its capacity to recognize and occupy inaccessible, nucleosomal motifs. We first performed spike-in normalized P3F ChIP-seq in Dbt/iP3F cells with 0 (t_0_), 8 (t_8_), and 24 (t_24_) hours of doxycycline treatment. At t_8_, we identified 28,740 high-confidence P3F binding sites enriched primarily with bZIP and bHLH motifs, while the PAX3:FKHR motif ranked 14^th^ among overrepresented sequences (**Figure 2A**). Interestingly, short bZIP DNA motifs share a high degree of sequence similarity with the central 5 nucleotides of the extended PAX3-FKHR motif in HOMER, suggesting this initial phase of P3F binding involves frequent sampling of partial motifs. By t_24_, as P3F protein expression plateaus (**Figure S4A**), the number of P3F sites was reduced to 9,552, with the PAX3:FKHR motif ranked 1^st^ among enriched sequences (**Figure 2A**). In addition to P3F binding sites being less numerous at t_24_ compared to t_8_, sites at t_24_ also had a lower average intensity (**Figures 2B** and **S4B**). These results may be consistent with the reported slow mobility of pioneer factors throughout chromatin as they sample non-specific nucleosomal sites before equilibration in a DNA sequence-driven manner (Lerner et al., 2020).

**Figure 2.**
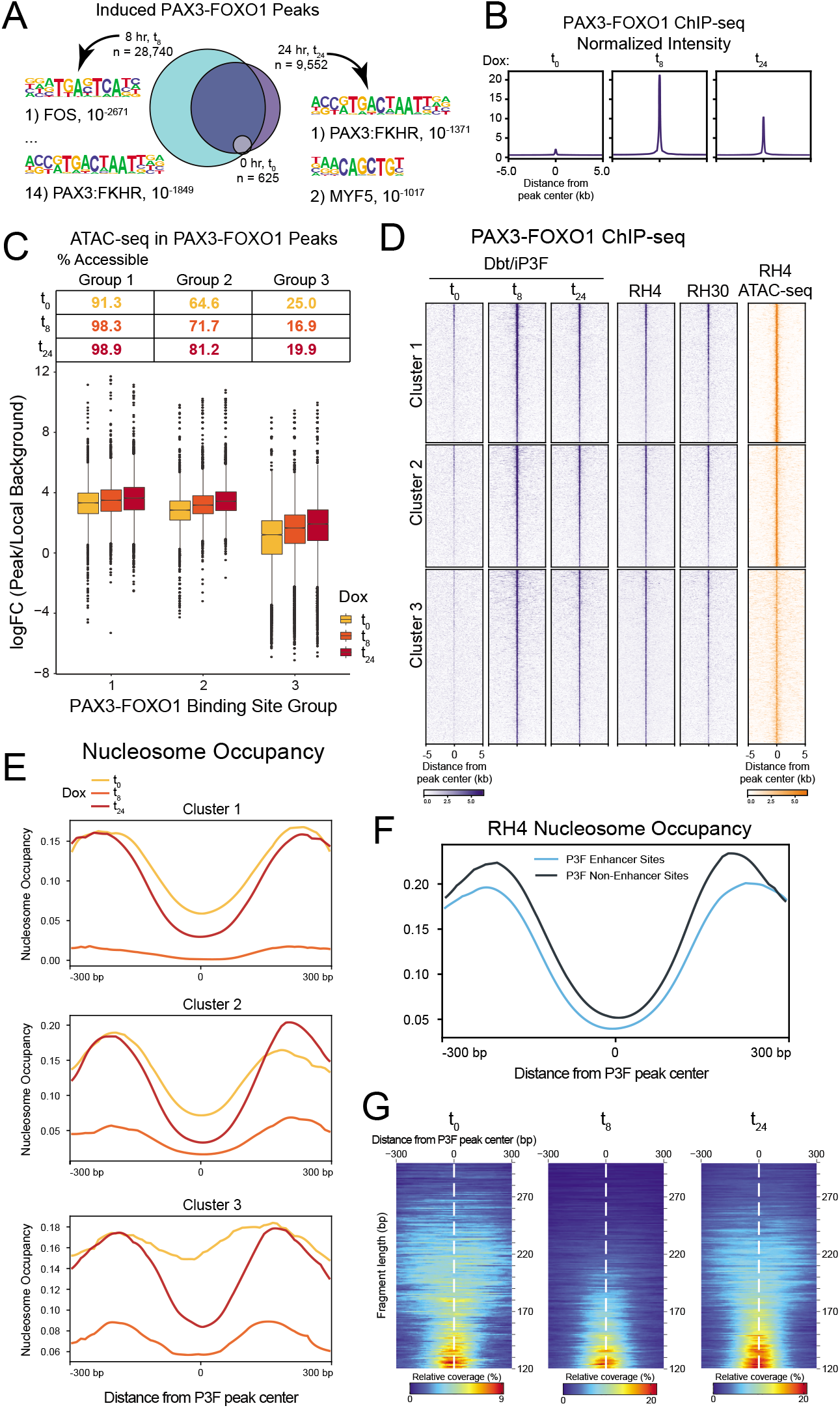
Early genomic targeting of PAX3-FOXO1 exhibits nucleosomal motif recognition. (**A**) PAX3-FOXO1 peaks called in Dbt/iP3F cells following treatment with 500 ng/mL doxycycline for 0 (t_0_), 8 (t_8_), or 24 (t_24_) hours. Motif analysis results are displayed for the t_8_ and t_24_ timepoints. (**B**) Spike-in normalized signal plot for PAX3-FOXO1 over the union of peaks identified at t_0_, t_8_, and t_24_. (**C**) ATAC-seq data from Dbt/iP3F cells treated with doxycycline for 0, 8, and 24 hours was utilized for k-means clustering (k = 3) of induced PAX3-FOXO1 binding sites. Local enrichment of ATAC-seq signals (calculated as the fold change of the ATAC signal within binding sites vs. 10X-sized background regions immediately flanking each binding site) is displayed for PAX3-FOXO1 binding site *Groups 1, 2,* and *3*. The table above the boxplots displays the percentage of sites within each *Group* that overlap ATAC-seq peaks at t_0_, t_8_, and t_24_. (**D**) Normalized ChIP-seq signal intensity and accessibility plotted across equilibrium PAX3-FOXO1 binding site *Clusters 1, 2,* and *3*. (**E**) NucleoATAC was performed using time course ATAC-seq data in Dbt/iP3F cells across equilibrium PAX3-FOXO1 binding site *Clusters 1, 2,* and *3*. Inferred nucleosome occupancy is displayed with respect to PAX3-FOXO1 peaks. (**F**) NucleoATAC of Enhancer and non-enhancer P3F binding sites in RH4 cells (**G**) plot2DO analysis of ATAC-seq fragment lengths present within induced PAX3-FOXO1 binding sites in Dbt/iP3F cells following 0, 8, and 24 hour doxycycline treatment.

ATAC-seq suggested that P3F sites in Dbt/iP3F cells universally exhibited increases in accessibility over the doxycycline treatment period, though we noted that many induced P3F sites begin with relatively high accessibility at t_0_ (**Figure S4C**). To understand if accessibility changes were limited to sites with low vs. high initial accessibility, we distinguished P3F binding sites in Dbt/iP3F cells according to their baseline accessibility signal. We defined P3F binding site *Groups 1, 2,* and *3* with high, medium, and low initial accessibility at t_0_, respectively (**Figure 2C**). In each *Group,* accessibility appeared to increase with time of P3F induction, though *Group 3* sites showed lower ATAC-seq signal at t_24_ than either *Group 1* or *2* sites at t_0_ (**Figure 2C**). In fact, a relatively small percentage of *Group 3* sites are defined as ATAC-seq peaks at t_0_, and few additional ATAC peaks overlapped *Group 3* sites at t_24_. We refined *Group 1, 2,* and *3* regions to include only P3F sites that are observed at equilibrium in RH4 cells (referred to as *Clusters 1, 2,* and *3*). ChIP-seq profiles in these regions indicate that the high P3F occupancy observed at t_8_ in Dbt/iP3F cells is a transient state prior to the cells approaching equilibrium at t_24_, more closely resembling the binding profiles observed in FP-RMS cells (**Figure 2D**). Additionally, we observed that the accessibility differences between *Clusters 1, 2,* and *3* in Dbt/iP3F cells were clearly visible in RH4 ATAC-seq data, revealing that the Dbt/iP3F model faithfully recapitulates the targeting and retention of P3F to relatively inaccessible regions (**Figures 2D** and **S4D**). Consistent with PF behavior reported previously, PAX3-FOXO1 binding in *Cluster 3* sites infrequently overlapped with ATAC-seq peaks at t_0_ or with induced ATAC-seq peaks at t_24_ (**Figure S4D**). Overall these patterns reveal that PAX3-FOXO1 rapidly targets to inaccessible chromatin regions, but the fusion protein is insufficient to induce accessibility at all binding locations. Differences in PAX3-FOXO1 signal intensity between 8- and 24-hour timepoints may reflect a late equilibrium period of P3F chromatin binding and release as additional factors are recruited to these regions.

Finally, we applied the NucleoATAC pipeline to determine if nucleosome positioning is affected over the time course of P3F induction and genomic occupancy (Schep et al., 2015). At t_0_ we observed strong evidence for nucleosome occupancy in *Cluster 1, 2,* and *3* sites (Figure S2E). Aligning nucleosome positions with respect to P3F peaks centers, we found that *Cluster 1* and *2* regions, exhibiting clear accessibility, had evidence of a pre-established nucleosome-depleted region (NDR) flanked by up- and downstream nucleosomes prior to P3F induction (Figure 2E). *Cluster 3* sites with low baseline accessibility, exhibited more uniform nucleosome occupancy prior to P3F induction, with less evidence of an NDR at t_0_. By t_24_ of P3F expression, the central NDR of *Clusters 1* and *2* appeared to be further remodeled, as evidenced by a widening and deepening of the nucleosome occupancy curve. Additionally, *Cluster 3* sites showed evidence of *de novo* establishment of an NDR directly overlapping a portion of induced P3F binding events while evidence for well-positioned nucleosomes neighboring these P3F peaks remained. This provides evidence of P3F targeting nucleosome-occupied sites followed in some cases by rapid remodeling events in inaccessible regions of the genome. We observed nearly identical patterns of equilibrium nucleosome occupancy in RH4 cells, suggesting that extended periods of P3F expression may ultimately result in the recruitment of additional factors required for nucleosome remodeling at many P3F sites (Figure 2F). We noted that NucleoATAC applied to our t_8_ ATAC-seq dataset was inefficient for inferring nucleosome positions in all *Clusters* (Figure 2E). Considering the NucleoATAC model inputs, we hypothesized that this was the result of differences in nucleosome-length fragments mapping to P3F sites, where a relative depletion of longer nucleosomal vs. shorter sub-nucleosomal fragments would result in low confidence nucleosome occupancy scores. Using plot2DO (Beati and Chereji, 2020), we in fact observed differences in fragment length distributions consistent with our NucleoATAC result (Figure 2G). We interpret this anomaly in nucleosome-length fragment distribution as evidence of local nucleosome redistribution taking place during the transition between baseline (t_0_) and P3F-driven epigenetic equilibrium (t_24_). Supporting the involvement of direct P3F binding in this nucleosome disruption event, we observed that Tn5 insertion rates generally increased over time in all P3F site *Clusters* (Figure S4F), and particularly in *Cluster 3,* we observed the gradual formation of a Tn5-protected footprint centered on high confidence PAX3:FKHR motifs at t_8_ and t_24_ (Figure S4G). While NucleoATAC provides some evidence that nucleosome positioning is altered upon P3F binding to chromatin, more than 70% of *Cluster 3* sites remain largely inaccessible even 24 hours after P3F expression (Figure S4D). This large subset of P3F binding events reveal that the fusion can establish a DNA occupancy footprint in regions otherwise inaccessible to traditional TFs, and is likely retained in these sites as necessary for the recruitment of additional chromatin remodeling factors.

## Conclusions

We have critically evaluated the PAX3-FOXO1 fusion oncoprotein with respect to categorical properties intrinsic to the pioneer class of transcription factors. Advances in recent years have catalyzed integrations across fields including pediatric oncology, genomics, and the biophysics of pioneer transcription factors (Fernandez Garcia et al., 2019; Nacev et al., 2020; Suva et al., 2013). Our studies have revealed that for two pioneer factors fused as a chimeric oncoprotein in a rare childhood tumor, chromatin recognition is consistent with PF function across the genome, including steady-state association with H3K9me3-marked heterochromatin domains and kinetic recognition of nucleosomal motifs. In developing quantitative <u>p</u>er-<u>c</u>ell normalization for our genome-wide binding studies (pc-ChIP-seq; **Methods**), we are able to infer nucleosome-targeting for PAX3-FOXO1, while accounting for sequencing bias resulting from the non-diploid genome structure of rhabdomyosarcoma models (Chen et al., 2015). We have discovered pioneer activity of the commonest driver alteration in fusion-positive rhabdomyosarcoma (Galili et al., 1993), which had previously uncharacterized function outside of active chromatin (Cao et al., 2010; Gryder et al., 2019; Gryder et al., 2017). Future efforts will be important to understand the kinetic rate constants of PAX3-FOXO1 dissociation from nucleosomes containing its motif (Donovan et al., 2019), as well as focused chromatin sequencing of sonication-resistant binding sites in heterochromatin (Becker et al., 2017). These efforts will be necessary to understand the role of PAX3-FOXO1 in heritable transmission of FP-RMS phenotypes through the establishment and maintenance of stable epigenetic states. In the coming years we anticipate continued convergence of fields, with emerging evidence to understand and predict the logic of pioneer activity in development and disease.

### Study Limitations

While our data provide strong evidence of PAX3-FOXO1’s pioneer function in FP-RMS cells, a number of limitations in our study encourage further investigation of this topic. More direct examination of nucleosomal motif binding is required. While the current study indicates that PAX3-FOXO1 targets nucleosome-occupied DNA rapidly upon induction, we rely on inferred nucleosome positioning computed from ATAC-seq data. While this method has proven reliable in reproducing chemically mapped nucleosome positioning in yeast (Schep et al., 2015), *in vitro* binding of purified PAX3-FOXO1 to nucleosome arrays assembled with DNA containing a PAX3-FOXO1 motif will provide a platform for measuring binding and dissociation rates. Genomic profiling of nucleosome positioning via MNase-seq will provide more direct evidence of nucleosomal binding and perturbation upon PAX3-FOXO1 expression.

Our study implies that the chromatin binding function of PAX3-FOXO1, to either accessible or repressed chromatin domains, may be insufficient to induce local chromatin activation, leading to context-dependent alteration in nearby gene expression. Assigning definitive gene regulatory roles to PAX3-FOXO1 in patient tumors is particularly challenging, given that we lack PAX3-FOXO1 genomic binding data in these samples. We are therefore faced with divergent hypotheses requiring experimental evidence: 1) PAX3-FOXO1 has a dominant chromatin structural role other than direct gene regulation (Rao et al., 2017) or 2) the logic of PAX3-FOXO1-dependent gene regulation involves additional requirements yet to be identified. This and other questions are brought to light by the discovery of a heterochromatin binding and nucleosome recognition activity of PAX3-FOXO1: opportunities to further our understanding of an oncogenic driver in a devastating pediatric sarcoma.

## Acknowledgements

We thank M.G. Poirier, M. Parthun, R.K. Wilson, C.E. Cottrell, B. Donovan, S.L. Lessnick, T.P. Cripe, C. Linardic, and all members of Stanton Lab for helpful discussion and comments. B.S. is supported by CancerFree KIDS (New Idea Award). F.G.B. is supported by the Intramural Research Program of the National Cancer Institute. B.Z.S. is grateful to support from the St. Baldrick’s Foundation (Berry Strong fund), Mark Foundation for Cancer Research (ASPIRE award), Andrew McDonough B+ Foundation (Childhood Cancer Research Grant).

## Author Contributions

Conceptualization, B.S. and B.Z.S.; Methodology, B.S., M.W., S.L., and B.Z.S.; Formal Analysis, B.S., M.W., and S.L.; Resources, F.G.B.; Data Curation, M.W., S.L., B.J.K., and J.R.F.; Writing – Original Draft, B.S., M.W., S.L., and B.Z.S.; Writing – Reviewing & Editing, B.J.K., J.R.F., F.G.B., P.W.; Visualization, B.S. and M.W.; Supervision, F.G.B., P.W., and B.Z.S.; Funding Acquisition, F.G.B. and B.Z.S.

## Supplemental Figures S1–S4

**Figure S1.**
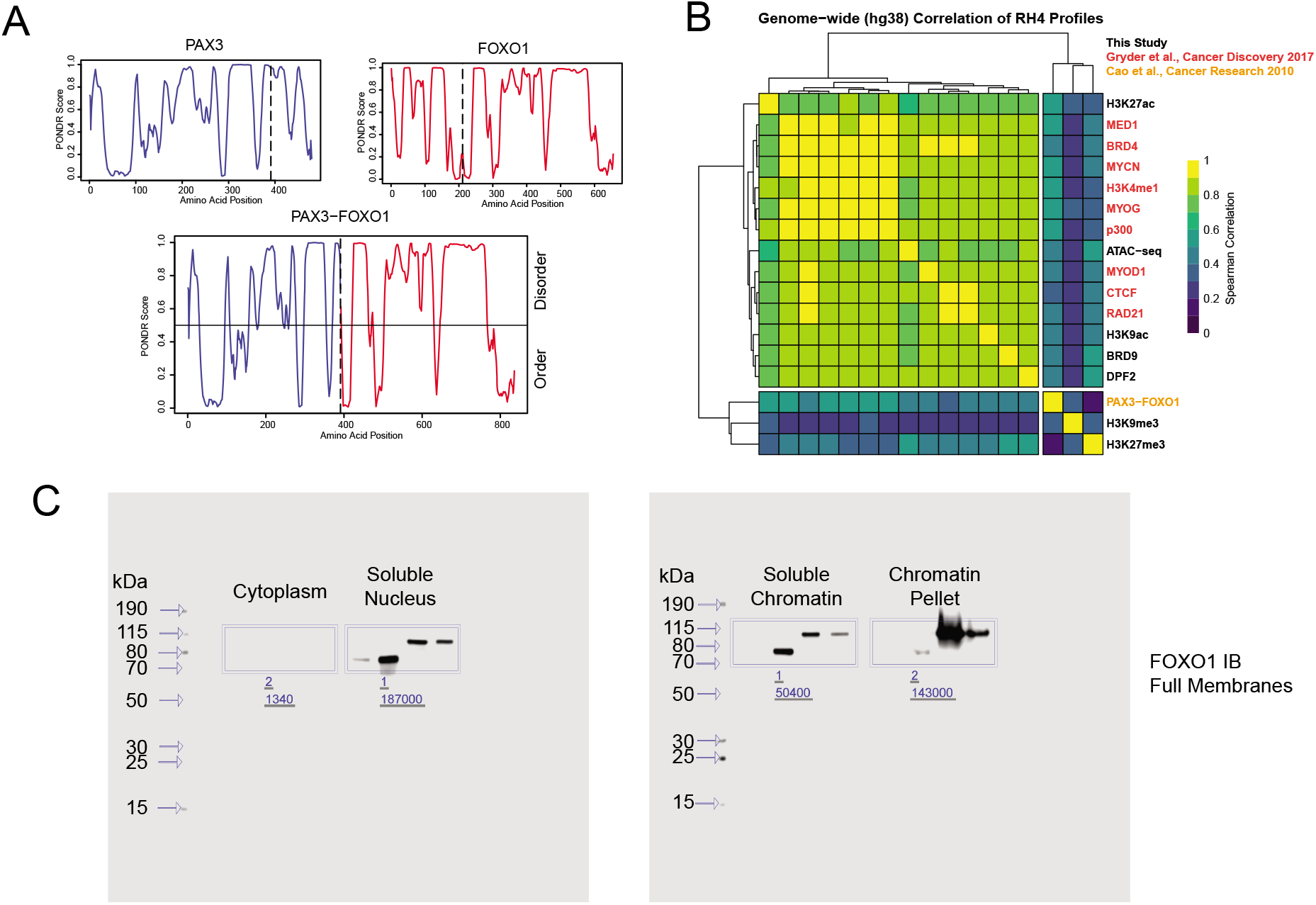
**(A)** PAX3, FOXO1, and PAX3-FOXO1 amino acid sequences were analyzed by PONDR to calculate regions of ordered and disordered protein structure. **(B)** Epigenomic profiles prepared in this study (H3K27me3, H3K27ac, H3K9ac, H3K9me3, DPF2, BRD9, and ATAC-seq) or available publicly (RAD21, CTCF, MYOD1, p300, MYOG, H3K4me1, MYCN, BRD4, MED1 – Gryder *et al., Cancer Discovery* 2017, and PAX3-FOXO1 – Cao *et al., Cancer Research* 2010) were used to compute genome-wide correlation values between each dataset. Clustered heatmap shows divergent binding profile for PAX3-FOXO1 with respect to euchromatic features profiled. **(C)** Full membranes of anti-FOXO1 western blots probing cellular fractions prepared from RMS cell lines displayed in **Figure 1B.**

**Figure S2.**
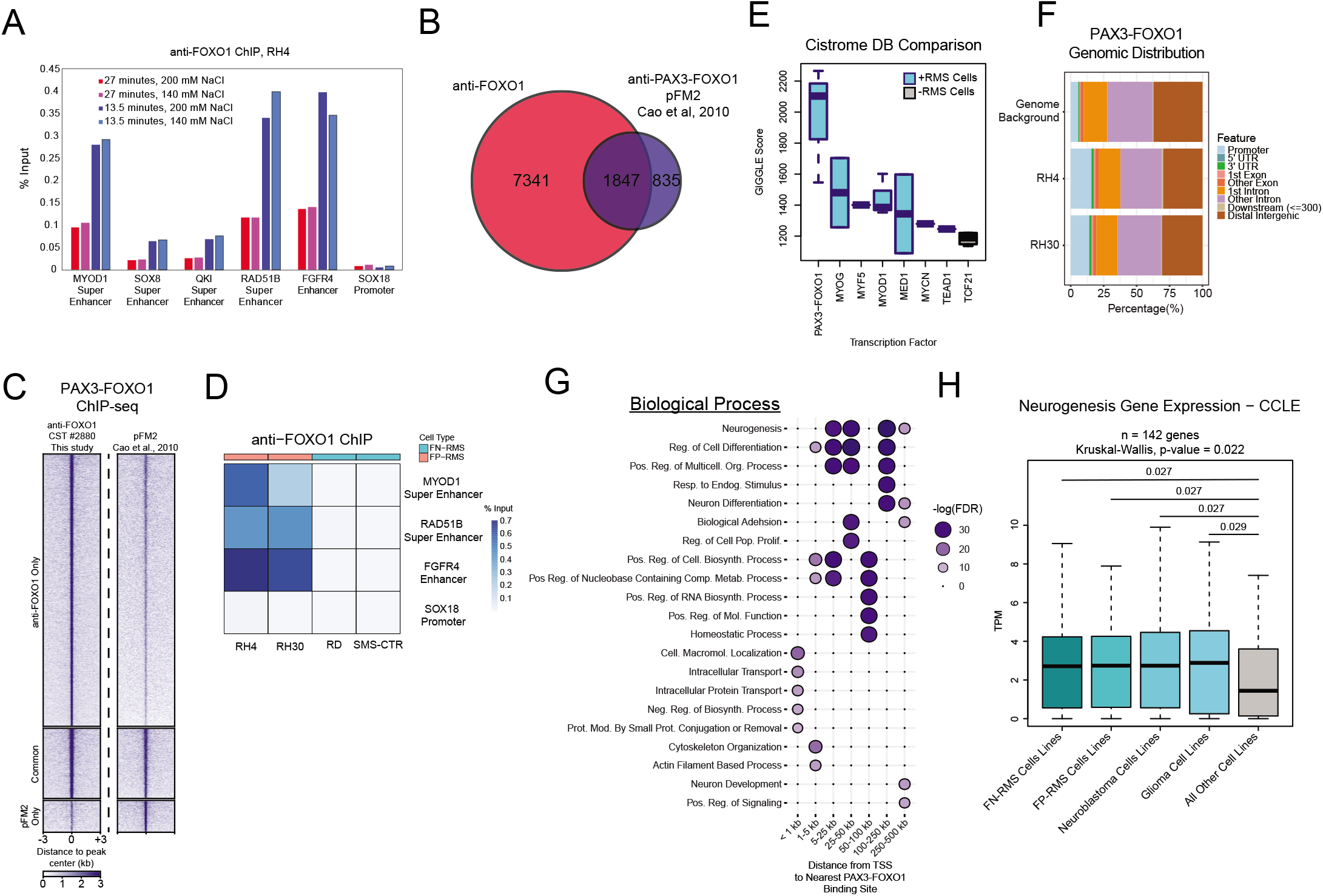
(**A**) ChIP-qPCR results following optimization of spike-in normalized PAX3-FOXO1 ChIP conditions in RH4 cells using a C-terminal FOXO1 antibody. Cells were sonicated for either 27 or 13.5 minutes and chromatin was immunoprecipitated in buffer containing either 200 or 140 mM NaCl. Optimal conditions were identified by comparing enrichment in known PAX3-FOXO1 binding sites (based on data from Cao *et al., Cancer Research* 2010) in the *MYOD1*, *SOX8, QKI, RAD51B,* and *FGFR4* loci compared to a negative control region in the *SOX18* promoter. (**B**) Overlap of PAX3-FOXO1 binding sites identified in this study with a FOXO1 antibody and sites identified by Cao *et al.* using an antibody (pFM2) spanning the PAX3-FOXO1 fusion breakpoint. (**C**) Heatmap of PAX3-FOXO1 enrichment in binding sites identified in this study, sites identified by Cao *et al.,* or both. (**D**) Ant-FOXO1 ChIP-qPCR results in a panel of FP-RMS and FN-RMS cell lines. (**E**) Comparison of PAX3-FOXO1 ChIP-seq in this study to publicly available ChIP-seq datasets using the Cistrome DB Toolkit. For factors profiled in at least one RMS cell line, boxplots are colored blue. Higher GIGGLE Score indicates stronger correlation between our PAX3-FOXO1 ChIP-seq data and the profiles for the indicated transcription factor. (**F**) Annotation of PAX3-FOXO1 binding sites identified in RH4 and RH30 cells. (**G**) PAX3-FOXO1 binding sites in RH4 cells were analyzed by GREAT. Genes linked to PAX3-FOXO1 binding sites were categorized according to distance from their TSS to the nearest binding site, and genes from each category were utilized for gene ontology analysis. The top 5 ontologies for each category are displayed. (**H**) 142 genes belonging to the gene ontology term “Neurogenesis” were queried against RNA-seq data from the Cancer Cell Line Encyclopedia. Expression level of these genes does not differ between FN-RMS, FP-RMS, Neuroblastoma, or Glioma cell lines, though these cell lines uniformly exhibit higher expression of neurogenic genes than all other cell lines (Kruskal-Wallis test, Benjamini-Hochberg p-value adjustment).

**Figure S3.**
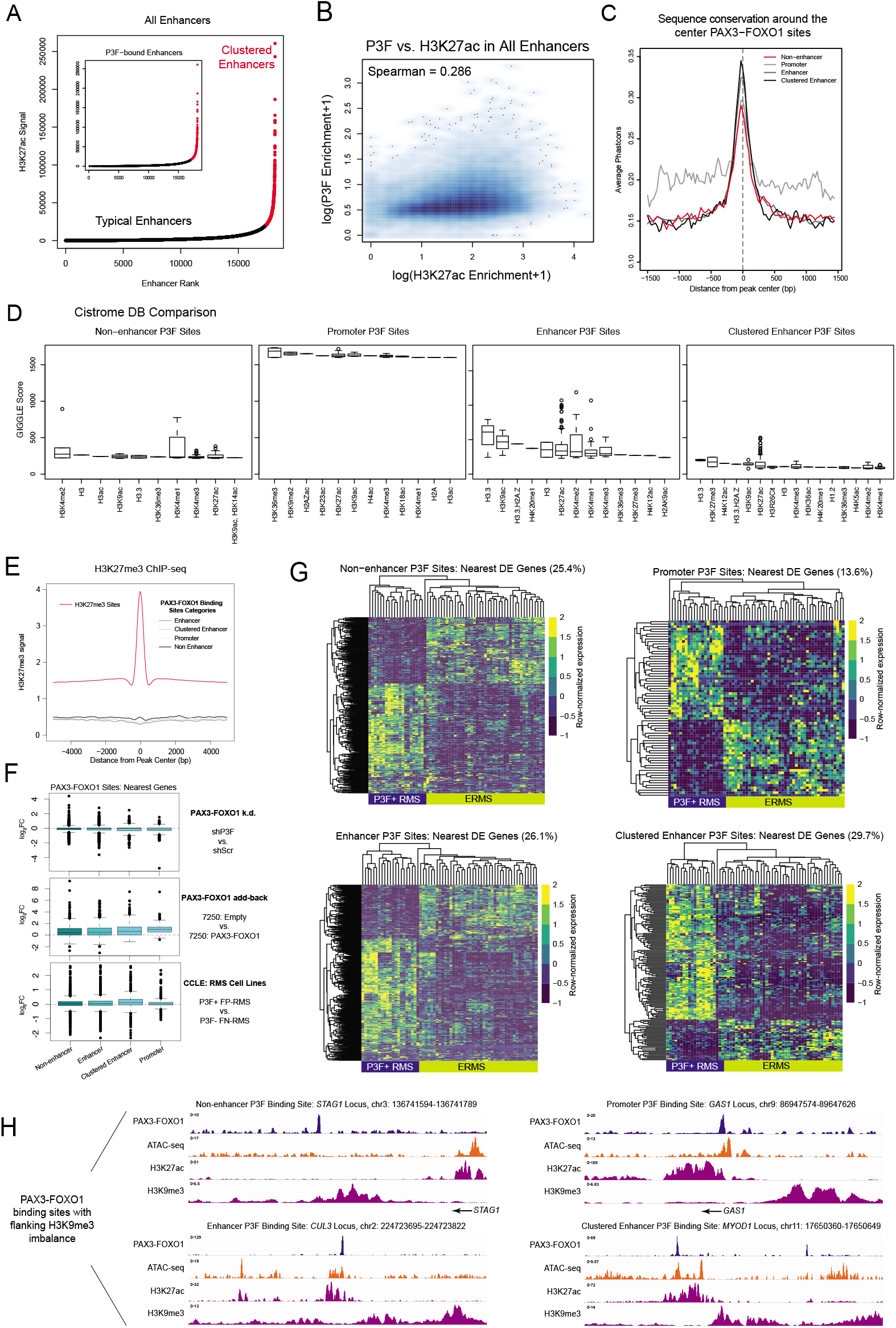
(**A**) H3K27ac ChIP-seq data was analyzed by ROSE to identify weaker, typical enhancers (black points) vs. strong, clustered enhancers (red points). Inset: P3F occupies enhancers across the entire spectrum of H3K27ac strength. (**B**) Comparison of P3F vs. H3K27ac signal enrichment in all enhancer regions. (**C**) Nucleotide sequence conservation of P3F binding site categories was calculated with phastCons. (**D**) The Cistrome DB Toolkit was used to assess similarity of publicly available histone modification/variant ChIP-seq profiles from various cell lines with P3F binding site categories in RH4 cells. High GIGGLE scores indicated stronger similarity. (**E**) H3K27me3 ChIP-seq profiles were generated for each P3F binding site category compared to true H3K27me3 peaks. (**F**) Analysis of public RNA-seq data. Genes linked to P3F binding site categories show little correlation with P3F expression. (**G**) RNA-seq data from the St. Jude Cloud Genomics Platform was analyzed to compare gene expression in P3F+ FP-RMS patient tumors compared to FN-RMS patient tumors. Differentially expressed (DE) genes (FC > 2, adjusted p-value < 0.05) associated with each P3F binding site category are displayed in a heatmap across all patient samples. Percentages represent the proportion of the total genes associated with each binding site category that show significant differential expression. (**H**) Example IGV plots of P3F binding sites in each category that exhibit asymmetric H3K9me3 enrichment. TSS position shown for *STAG1* and *GAS1. CUL3* and *MYOD1* TSS is outside of the viewer window. Chromosomal coordinates indicate P3F binding site location.

**Figure S4.**
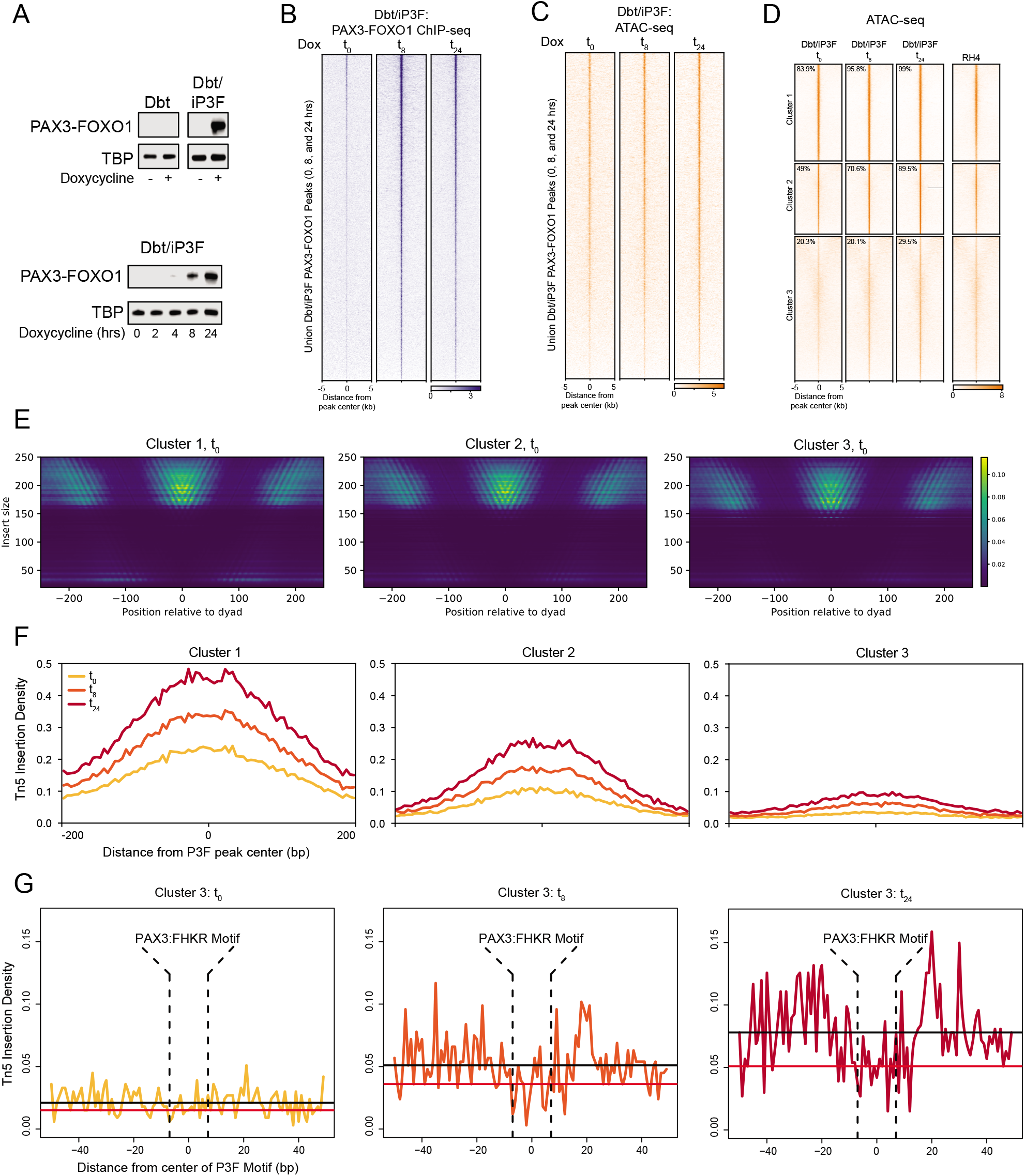
(**A**) (Top) Western blot analysis of PAX3-FOXO1 protein expression in parental Dbt and inducible Dbt/iP3F cells following 24-hour treatment with 500 ng/mL doxycycline. (Bottom) Western blot analysis of PAX3-FOXO1 induction in Dbt/iP3F cells treated with 500 ng/mL doxycycline for the indicated time. (**B**) Heatmap of spike-in normalized PAX3-FOXO1 pc-ChIP-seq in Dbt/iP3F cells following 0 (t_0_), 8 (t_8_), or 24 (t_24_) hours of doxycycline treatment. (**C**) Heatmap of ATAC-seq in Dbt/iP3F cells following 0, 8, or 24 hours of doxycycline treatment. Vertical arrangement of peaks in **B** and **C** are independent. (**D**) Dbt/iP3F (t_0_, t_8_, and t_24_) and RH4 ATAC-seq heatmaps over equilibrium PAX3-FOXO1 binding site *Clusters 1, 2,* and *3*. Percentages represent the number of P3F binding sites in each *Cluster* that overlap with ATAC-seq peaks at each timepoint. (**E**) NucleoATAC analysis of baseline (t_0_) nucleosome positioning across equilibrium PAX3-FOXO1 binding site *Clusters* in Dbt/iP3F cells. Nucleosome- and sub-nucleosome-length ATAC-seq fragments are plotted with respect to high confidence nucleosome positions. (**F**) Tn5 insertion patterns from NucleoATAC were plotted over equilibrium PAX3-FOXO1 binding site *Clusters.* (**G**) Tn5 insertions were plotted with respect to high confidence PAX3:FKHR motif positions (indicated by vertical dashed lines) within *Cluster 3* binding sites. Tn5 insertion patterns reveal formation of a protected PAX3-FOXO1 footprint over time from t_0_ (left) to t_24_ (right) of doxycycline treatment. The median insertion rate outside and within the motif positions is displayed as a black and red horizontal line, respectively.

### Methods

#### RESOURCE AVAILABILITY

##### Lead Contact

Further information and requests for resources and reagents should be directed to and will be fulfilled by the Lead Contact, Dr. Benjamin Stanton.

##### Materials Availability

This study did not generate new unique reagents.

##### Data and Code Availability

Raw sequence files for each dataset prepared for this manuscript are publicly available through the Gene Expression Omnibus (GEO, GSE163068). Also available are processed files (hg38) generated for immediate use. These include all genome-wide signal files (bigwig format) and PAX3-FOXO1 peak calls (bed format). PAX3-FOXO1 signals and peaks are directly comparable across cell lines and conditions, as they have been collectively normalized for this study. Clinical RNA-seq data for pediatric rhabdomyosarcoma patients used for analysis in this study were obtained from the St. Jude Cloud (McLeod et al., 2021). Only publicly available algorithms were used in the analysis of datasets presented in this study. Scripts will be made available upon request.

#### EXPERIMENTAL MODEL AND SUBJECT DETAIL

##### Cell Lines

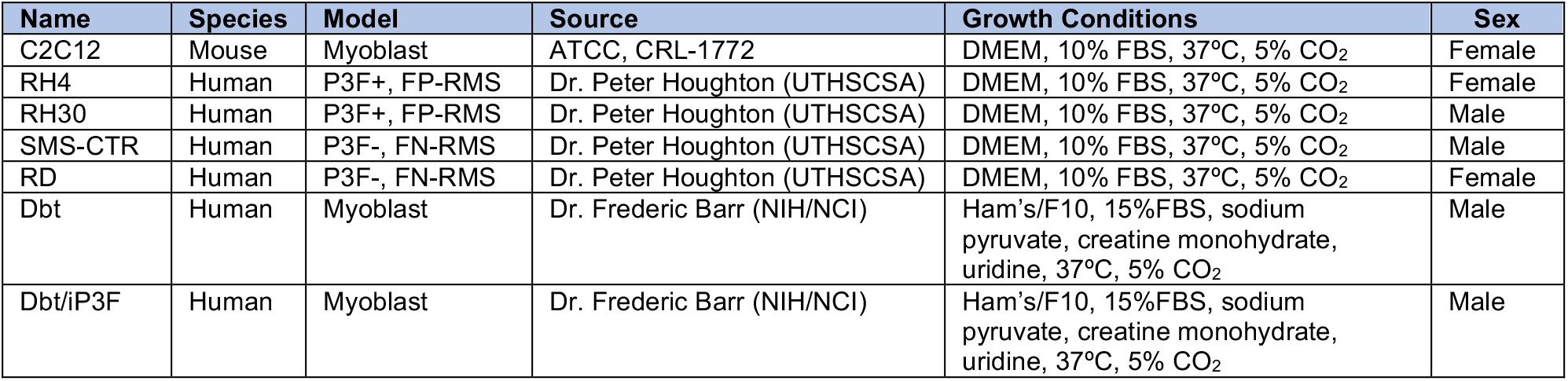

| Name | Species | Model | Source | Growth Conditions | Sex |
| --- | --- | --- | --- | --- | --- |
| C2C12 | Mouse | Myoblast | ATCC, CRL-1772 | DMEM, 10% FBS, 37°C, 5% CO<sub>2</sub> | Female |
| RH4 | Human | P3F+, FP-RMS | Dr. Peter Houghton (UTHSCSA) | DMEM, 10% FBS, 37°C, 5% CO<sub>2</sub> | Female |
| RH30 | Human | P3F+, FP-RMS | Dr. Peter Houghton (UTHSCSA) | DMEM, 10% FBS, 37°C, 5% CO<sub>2</sub> | Male |
| SMS-CTR | Human | P3F-, FN-RMS | Dr. Peter Houghton (UTHSCSA) | DMEM, 10% FBS, 37°C, 5% CO<sub>2</sub> | Male |
| RD | Human | P3F-, FN-RMS | Dr. Peter Houghton (UTHSCSA) | DMEM, 10% FBS, 37°C, 5% CO<sub>2</sub> | Female |
| Dbt | Human | Myoblast | Dr. Frederic Barr (NIH/NCI) | Ham's/F10, 15%FBS, sodium pyruvate, creatine monohydrate, uridine, 37°C, 5% CO<sub>2</sub> | Male |
| Dbt/iP3F | Human | Myoblast | Dr. Frederic Barr (NIH/NCI) | Ham's/F10, 15%FBS, sodium pyruvate, creatine monohydrate, uridine, 37°C, 5% CO<sub>2</sub> | Male |

#### METHOD DETAILS

##### Cell Culture

RH4 (FP-RMS), RH30 (FP-RMS), RD (FN-RMS), and SMS-CTR (FN-RMS) cells, a gift from Dr. Peter Houghton (UTHSCSA), were cultured in high-glucose DMEM supplemented with 10% FBS, Glutamax, and penicillin/streptomycin. Immortalized Dbt myoblasts engineered with doxycycline-inducible PAX3-FOXO1 (Dbt/iP3F), engineered in the lab of Dr. Frederic Barr (NIH/NCI) (Pandey et al., 2017), were cultured in Ham’s/F-10 supplemented with 15% FBS, glutamine, sodium pyruvate, creatine monohydrate, uridine, and penicillin/streptomycin. Mouse C2C12 myoblasts were purchased from ATCC and cultured in high-glucose DMEM supplemented with 10% FBS, Glutamax, and penicillin/streptomycin. For PAX3-FOXO1 induction studies, Dbt/iP3F cells were seeded in normal culture media and grown to approximately 70% confluence before exchange with media containing 500 ng/mL doxycycline hyclate for the desired time period (8 or 24 hrs).

##### Cell Fractionation and Immunoblotting

Cells were collected from 15 cm plates and washed with ice-cold PBS containing protease inhibitor cocktail. Cell pellets were resuspended in Fractionation Buffer 1 (20 mM HEPES, 10 mM KCl, 0.2 mM EDTA) with protease inhibitor cocktail and incubated for 10 minutes on ice. NP-40 was added to a final concentration of 0.5% and the samples was vortexed on high for 15 seconds. Samples were incubated on ice for 1 minute, vortexed at high speed for 15 seconds, and pelleted at 14,000 rpm for 1 minutes at 4°C. The supernatant (cytoplasmic fraction) was transferred to a clean Eppendorf tube, and the nuclei pellet was resuspended in Fractionation Buffer 2 (10 mM Tris-HCl pH 8.0, 1 mM EDTA, 0.1% NP-40, 500 mM NaCl) with protease inhibitor cocktail and incubated at 4°C for 45-60 minutes with overhead rotation. The sample was centrifuged at 14,000 rpm for 10 minutes and the supernatant (soluble nuclear fraction) was transferred to a clean Eppendorf tube. The remaining chromatin pellet was resuspended in IP Buffer (50 mM Tris-HCl pH 8.0, 300 mM NaCl, 1 mM EDTA, 1% Triton X-100) with protease inhibitor cocktail and sonicated with an Active Motif EpiShear probe sonicator equipped with a cooled sonication platform for 5 minutes at 30% amplitude cycling from 30 seconds ON to 30 seconds OFF. The sample was centrifuged at 14,000 rpm for 10 minutes at 4°C and the supernatant (soluble chromatin fraction) was transferred to a clean Eppendorf tube. The remaining chromatin pellet was resuspended in 2X SDS protein sample buffer with 10% v/v 2-mercaptoethanol and heated at 95°C for 10 minutes. The other cell fractions were quantified with the Pierce Rapid Gold BCA Protein Assay Kit. Samples were resolved by SDS-PAGE using 2.5 μg of the cytoplasmic, soluble nuclear, and soluble chromatin fractions or 10 μL of the chromatin pellet fractions on a NuPAGE 4-12% Bis-Tris gel.

Proteins were transferred overnight at 4°C/30V to nitrocellulose membranes. Membranes were blocked at room temperature in 5% w/v milk solution in TBST (0.1% v/v Tween-20) for 1 hour before incubation for 2 hours at room temperature with primary antibodies detecting: MYCN (Cell Signlaing, 51705S), BAF155 (Cell Signaling, 11956S), FOXO1 (Cell Signaling, 2880S), HP1α (Cell Signaling, 2616S), H3K9me3 (Active Motif, 39062), H4K20me1 (Active Motif, 39727), or TBP (Cell Signaling, 44059S). Following 1 hour incubation with HRP-linked secondary antibodies, immunoblots were incubated with SuperSignal West Pico PLUS Chemiluminescent Substrate, images were acquired with a LI-COR C-DiGit Blot Scanner running Image Studio v5.2.

##### Amino Acid Primary Sequence Analysis

Human PAX3 (Uniprot ID: P23760) and FOXO1 (Uniprot ID: Q12778) amino acid sequence conservation was analyzed on the ConSurf Server using default parameters (Berezin et al., 2004). ConSurf output files were used to project evolutionary conservation estimates, buried/exposed residue classifiers, and functional/structural residue classifiers onto the primary protein structure in **Fig. 1A**. Protein disorder analysis was conducted using the VL-XT PONDR algorithm (Romero et al., 2001) on the wild type PAX3 and FOXO1 protein sequences as well as the PAX3-FOXO1 fusion protein sequence.

##### PAX3-FOXO1 Spike-in ChIP Optimization

To facilitate quantitative normalization across ChIP-seq samples not only from different treatment conditions but also from distinct cell lines, we employed a spike-in ChIP strategy using known numbers of cells of human origin mixed in defined ratios with cells of mouse origin. This approach is similar in theory and in practice to the recently reported quantitative HiChIP method, known as AQuA-HiChIP (Gryder et al., 2020). Our spike-in strategy addresses technical confounders introduced at two key points in the ChIP-seq protocol. Firstly, we introduce formaldehyde fixed mouse C2C12 cells to fixed human RMS or myoblast cells prior to sonication. In this study, the ratio of human:mouse cells is fixed at 3:1 for all experiments. This strategy ensures subsequent normalization steps retain read depth information on the basis of starting cell number. Failure to do so may obscure differences in chromatin output per cell across different cell types and treatment conditions. The assumption of equal chromatin produced per cell by sonication of any cell type implicit in commercial spike-in normalization reagents may be frequently violated when conducting experiments comparing aneuploid cancer cell lines, or in our case, RMS cells in which genome duplication may be a frequent event. Secondly, as in previous ChIP-Rx and commercial spike-in strategies (Egan et al., 2016; Orlando et al., 2014), we assume an equal number of mouse DNA fragments comprise the final sequencing libraries for input samples and for ChIP samples that will be directly compared. This permits us to properly correct for differences in sequencing depth across samples based on an internal and constant reference.

Our strategy utilizes one antibody, which we ideally expect to react with specific, conserved epitopes on human and mouse chromatin (e.g. histone modifications). In this ideal case, reliable and stable ChIP efficiency against epitopes on mouse chromatin ensures adequate and reproducible read numbers mapping to the spike-in mouse genome across all samples, while distinct human cell lines (e.g. RMS cells vs. myoblasts) under various treatment conditions (e.g. doxycycline induction) may produce a variable number of reads mapping to the human genome. In the case that an antibody has exquisite species-specific reactivity, or the desired epitope is not expressed in C2C12 cells (e.g. lineage restricted transcription factors), we leverage the inherently low signal-to-noise ratio of ChIP assays to our advantage. Here the number of non-specific reads mapping to the mouse genome is anticipated to remain constant across all ChIP samples, as even in highly efficient ChIP assays, background reads are present in relatively high proportions. Therefore, a single antibody approach is sufficient to produce ChIP samples with a constant number of spike-in reads mapping to the mouse genome for a given antibody, regardless of the reactivity of that antibody with mouse epitopes.

In addition to developing a novel, per-cell ChIP (pc-ChIP) approach, we tested two variables to optimize conditions for PAX3-FOXO1 ChIP using a FOXO1 antibody: 1) sonication time and 2) salt concentration in the ChIP buffer. The detailed pc-ChIP optimization strategy follows:

###### Cell fixation

FP-RMS, FN-RMS, Dbt/iP3F, or C2C12 cells were cultured in 15 cm plates, dissociated with trypsin, pelleted, and washed with PBS. Cell pellets were resuspended in Fixing Buffer (50 mM HEPES pH 7.3, 1 mM EDTA, 0.5 mM EDTA, 100 mM NaCl) and fresh, methanol-free formaldehyde was added to a final concentration of 1%. After 10 minutes of incubation at room temperature, the fixation was quenched by the addition of glycine to a final concentration of 125 mM and the cell suspension was placed on ice for 5 minutes. Fixed cells were pelleted at 1,200 x g for 5 minutes at 4°C and resuspended in ice-cold PBS containing a protease inhibitor cocktail. Fixed FP-RMS, FN-RMS, and Dbt/iP3F cells were aliquoted at 6×10^6^ cells/tube, and fixed C2C12 cells were aliquoted at 2×10^6^ cells/tube. Cells were pelleted at 3,000 rpm for 5 minutes at 4°C, the supernatant was removed, and pellets were snap frozen and stored at −80°C until further use.

###### Sonication

For initial optimization in RH4 cells, thawed aliquots of 6×10^6^ RH4 cells were combined with thawed aliquots of 2×10^6^ C2C12 cells in 800 uL of TE buffer pH 8.0 containing protease inhibitor cocktail and transferred to polystyrene sonication tubes. Samples were sonicated with an Active Motif EpiShear probe sonicator equipped with a cooled sonication platform for either 27 minutes or 13.5 minutes at 30% amplitude cycling from 30 seconds ON to 30 seconds OFF. After sonication, a 5 uL volume was aliquoted from each sample (input) and combined with 20 uL TE, 1 uL 10% SDS, and 1 uL 20 mg/mL Proteinase K for overnight decrosslinking at 65°C. The remaining sonicated chromatin was stored at 4°C. Input samples were purified using Qiagen MinElute PCR Purification columns and chromatin fragmentation was assessed by gel electrophoresis on an E-Gel 2% EX agarose gel.

###### ChIP

After evaluating fragmentation, sonicated chromatin in TE buffer was adjusted to ChIP Buffer by the addition of Triton X-100 (to 1% final concentration), SDS (to 0.1% final concentration), and sodium deoxycholate (to 0.1% final concentration). For initial optimization, we added sodium chloride to a final concentration of either 140 mM or 200 mM. Chromatin in ChIP Buffer was incubated on ice for 5 minutes and insoluble material was removed by centrifuging the sample at 13,000 rpm for 10 minutes at 4°C and transferring the supernatant to a new 1.5 mL tube. 5 uL of FOXO1 antibody (Cell Signaling Technology, #2880) was added, and samples were incubated at 4°C for 1 hour with overhead rotation. For each sample, 40 uL of Protein A Dynabeads were buffer exchanged with ChIP buffer containing either 140 mM or 200 mM NaCl before addition to the antibody:chromatin mixture, and the samples were incubated overnight at 4°C with overhead rotation. The beads were then washed twice with Low Salt Wash Buffer (0.1% SDS, 0.1% sodium deoxycholate, and 1% Triton X-100 in TE buffer pH 8.0), twice with ChIP Buffer (containing either 140 mM or 200 mM NaCl), twice with LiCl Wash Buffer (250 mM LiCl, 0.5% NP-40, 0.5% sodium deoxycholate in TE buffer pH 8.0), and twice with TE buffer pH 8.0. The beads were then resuspended in 100 uL TE buffer pH 8.0 with 2.5 uL 10% SDS and 5 uL 20 mg/mL Proteinase K, and ChIP samples were decrosslinked at 65°C overnight. ChIP DNA was purified with Qiagen MinElute PCR Purification columns.

###### Real-time PCR validation

Efficiency and specificity of our PAX3-FOXO1 ChIP conditions were first assessed by real-time PCR with primers designed against known PAX3-FOXO1 binding sites within the *MYOD1, SOX8, QKI, RAD51B,* and *FGFR4* loci (primer sequences listed below) (Cao et al., 2010). A negative control region in the *SOX18* promoter was also tested (Yohe et al., 2018). This analysis revealed specific and robust ChIP enrichment within known PAX3-FOXO1 binding sites compared to the negative control region.

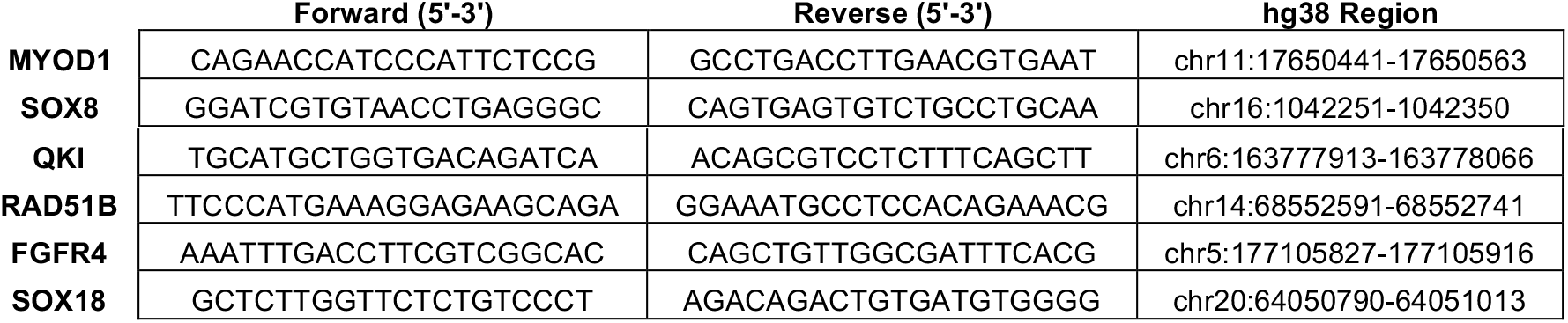

This optimization strategy revealed that 13.5 minutes of sonication followed by ChIP in buffer containing 200 mM NaCl was suitable for robust PAX3-FOXO1 ChIP enrichment using a FOXO1 antibody with limited background signal. All subsequent PAX3-FOXO1 ChIP assays in RH4, RH30, RD, SMS-CTR, and Dbt/iP3F cells were performed in an identical fashion.

Additional ChIP assays performed for this study were conducted in RH4 cells (with C2C12 spike-in) sonicated for 27 minutes, with immunoprecipitation performed in ChIP buffer with 200 mM NaCl using antibodies against DPF2 (Abcam, ab134942), BRD9 (Abcam, ab137245), H3K27ac (Active Motif, 39133), and H3K27me3 (Active Motif, 39155). H3K9me3 (Active Motif, 39062) and H3K9ac (EpiCypher, 13-0020) ChIPs were performed in RH4 cells (with C2C12 spike-in) sonicated for 12 minutes.

##### pc-ChIP-seq Considerations

We perform our pc-ChIP-seq spike-in and normalization on a “per cell” basis to retain information regarding the chromatin output across cell lines tested under various treatment conditions. Conversely, commercially available spike-in reagents recommend an equal starting amount of chromatin for each sample, regardless of starting cell number. For direct, quantitative comparison to be performed between distinct cell lines and treatments, the condition of equal chromatin produced per cell in all groups must be satisfied when normalizing on the basis of starting chromatin amount. In RMS cell lines and other aneuploid cancer cell lines, this condition is grossly violated. These cell lines generally contain variably sized genomes, and therefore they release different amounts of chromatin per cell that can globally influence the signal output from a standard ChIP assay. Using human/mouse read ratios in our input sequencing libraries, we can demonstrate the need to account for this by inferring the relative genome size for each cell line in this study. We used tetraploid C2C12 mouse cells for spikein (estimated genome size, 5.4×10^9^ bp) at a 3 to 1 ratio of human to mouse cells. We can therefore calculate the inferred genome sizes as:

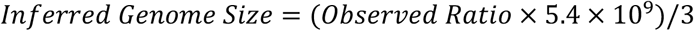

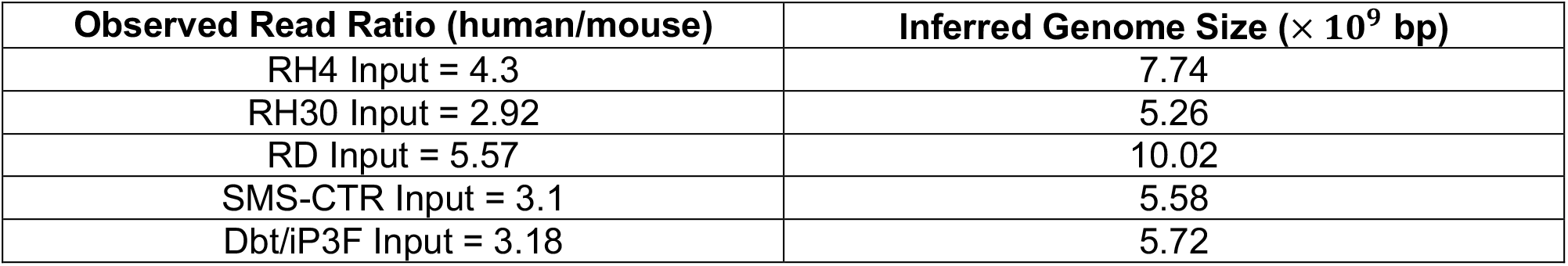

These figures reveal vastly different relative genome sizes across cell lines tested and showed consistent results across RH4 replicates (7.72 and 7.92 x10^9^ bp, respectively). This is consistent with the scenario where chromatin output differs globally, and therefore relative read depth across both input and ChIP samples differ. Under these conditions, a spike-in and normalization procedure based on the starting amount of *chromatin* instead of the starting number of *cells* would obscure the differences between different biological samples. This prevents qualitative comparison of peak calls as wells as quantitative comparison of signal strength observed. Therefore, we needed a new strategy where the difference of amount of chromatin released by each cell is taken into consideration, and we developed pc-ChIP-seq to correct this bias. While the focus here is on chromosome ploidy resulting in differential chromatin output per cell, epigenetic repression and de-repression as well as cell cycle stage are examples of other conditions that may globally influence chromatin output per cell owing to differences in sonication sensitivity.

To justify the assumption that a one-antibody strategy is sufficient for a spike-in ChIP-seq normalization approach, even in the event that the chosen antibody does not recognize epitopes on mouse chromatin to produce high quality ChIP-seq data in the spike-in genome, we first analyzed our anti-FOXO1 ChIP-seq data from RH4/C2C12 cells with the ENCODE pipeline, aligning only to mm10. This analysis, prior to down-sampling, revealed just 135 low-confidence FOXO1 peaks in the mouse genome. More importantly, just 0.18% of all reads aligning to the mouse genome mapped to these peaks. Second, in replicate PAX3-FOXO1 ChIP-seq datasets produced in RH4 cells, the ratio of human to mouse reads in the ChIP samples were 4.57 and 4.52. Together, these findings suggest even when non-specific mouse reads are not informative for peak calling in the mouse genome, they comprise a substantial and reproducible portion of a ChIP sample that can be leveraged for normalization. This reflects the inherently low signal-to-noise ratio of ChIP assays, where a typical transcription factor ChIP will often have a relatively low Fraction of Reads in Peaks (FRiP) value.

##### pc-ChIP Library Construction and Sequencing

ChIP and input DNA from each sample were prepared for sequencing as before (Kidder and Zhao, 2014) by blunt end repair using the Lucigen End-It DNA End-Repair Kit, 3’ A-tailing by Klenow fragment (3’-5’ exo-), adaptor ligation by T4 DNA ligase, and size selection on an E-Gel 2% EX agarose gel. Libraries were amplified with barcoded primers and isolated from unreacted primers by gel purification. H3K9me3 ChIP and Input DNA samples were sonicated for 2 minutes (30 seconds ON/30 seconds OFF) prior to library preparation, as in (Becker et al., 2017). Following adaptor ligation and PCR amplification steps, unreacted adaptors/primers were removed with AMPure XP beads (1.8:1 ratio), and no further library size-selection was performed. Pooled libraries were sequenced at the Nationwide Children’s Hospital Institute for Genomic Medicine, Genomic Services Laboratory on a HiSeq4000 running in paired-end, 150 bp mode.

##### pc-ChIP-seq Normalization

As the pc-ChIP protocol rigorously controls for the starting number of cells and the ratio of human to mouse cells across all samples in each experimental group, read depth bias correction is achieved through normalization using mouse spike-in reads across all input and ChIP samples for a given comparison. We carried out normalization prior to peak calling and generation of signal files in order to perform quantitative comparisons across cell lines or test samples. In detail, our approach for random down-sampling of sequencing reads across samples is as follows:

BCL converted paired end fastq files were aligned to hg38 and mm10 reference genomes, separately, utilizing bowtie2 (Langmead and Salzberg, 2012) and following ENCODE best practices. The resulting mouse BAMs were analyzed by samtools to calculate the number of reads aligned to each genome. The minimum number of mouse aligned reads (*m*) was identified across all samples, and was divided by the number of aligned mouse reads for each sample (*s*) to calculate the scaling factor (*f*), *f=m/s*. This scaling factor was subsequently used to normalize the hg38 aligned BAMs through subsampling with samtools view -s, which retains read pair information. Picard SamToFastq converted the resulting SAM files to paired end fastq files.

##### ChIP-seq Data Analysis

Normalized pc-ChIP-seq fastq files from this study as well as publicly available PAX3-FOXO1 ChIP-seq fastq files for RH4 cells (GSE19063, (Cao et al., 2010)) were processed with the ENCODE ChIP-seq pipeline (v1.3.4) with chip.xcor_exclusion_range_max set at 25. Normalized, paired-end fastqs were aligned with bowtie2 (version 2.3.4.3) to hg38, with parameters bowtie2 -X2000 -mm. Next, blacklisted region, unmapped, mate unmapped, not primary alignment, multi-mapped, low mapping quality (MAPQ<30), duplicate reads and PCR duplicates were removed. Peaks were called with MACS2 (version 2.2.4), with parameters -p 1e-2 --nomodel --shift 0 -- extsize $[FRAGLEN] --keep-dup all -B -SPMR, where FRAGLEN is the estimated fragment length. IDR analyses were performed on peaks from replicate samples or pseudo-replicates, with threshold 0.05. Motif analysis with HOMER (version 4.11.1) (Heinz et al., 2010) was then carried out on conservative IDR peaks. For visualization, bedGraph files were generated with MACS2 bdgcmp from the pile-up, and then converted to bigwig format with bedGraphToBigWig. Heatmaps were generated with deeptools (version 3.3.1). k-means classification from deeptools was used to generate peak clusters for **Fig. 2**. ROSE (v0.1) (Whyte et al., 2013) was used with our H3K27ac IDR peaks to classify enhancers and to stitch proximate H3K27ac regions into clustered enhancers.

##### ATAC-seq and ATAC-seq Data Analysis

ATAC-seq was performed as previously described (Buenrostro et al., 2013) with only minor modifications. 5×10^4^ cells per experiment were first washed with RSB buffer (10 mM Tris-HCl pH 8, 10 mM NaCl, 3 mM MgCl_2_) and gently permeabilized with RSB lysis buffer (10 mM Tris-HCl pH 8, 10 mM NaCl, 3 mM MgCl_2_, 0.1% NP-40) on ice. Cells were suspended in 50 uL of tagmentation master mix prepared from Illumina Tagment DNA TDE1 Enzyme and Buffer Kit components (#20034197), and transposition was performed for 30 minutes at 37°C. Tagmented DNA fragments were isolated using Qiagen MinElute PCR Purification columns prior to library amplification. ATAC-seq libraries were amplified with barcoded Nextera primers for 14 cycles, and excess primers were removed by size selection with AMPure XP beads. Libraries were sequenced on the HiSeq4000 platform running in PEx150bp mode.

The ENCODE ATAC-seq pipeline (https://github.com/ENCODE-DCC/atac-seq-pipeline) with default parameters was used to process ATAC-seq data. First, reads are scanned for adaptor sequences and trimmed with cutadapt (version 2.3). Reads are then mapped to hg38 with bowtie2 (version 2.3.4.3). Properly aligned, non-mitochondrial read pairs were retained for peak calling with MACS2 (version 2.2.4). After peaks are called, heatmaps are generated with deeptools (version 3.3.1) (Ramirez et al., 2016). Fragment length distributions were generated with plot2DO v1.0 (https://github.com/rchereji/plot2DO) (Beati and Chereji, 2020). Local signal vs background enrichment is calculated with localEnrichmentBed (https://github.com/dariober/bioinformatics-cafe/tree/master/localEnrichmentBed). Nucleosome position, nucleosome occupancy and tn5 insertion density are estimated using NucleoATAC (version 0.3.4) (Schep et al., 2015). Transcription factor foot printing/motif protection was assessed by identifying PAX3-FKHR or FOXO1 motif positions within PAX3-FOXO1 binding sites using FIMO (Grant et al., 2011). Insertion rates were subsequently plotted over aligned motif positions using deeptools (version 3.3.1).

##### RNA-seq Analysis

Tumor tissue expression data (in bam format) from pediatric patients diagnosed with Alveolar Rhabdomyosarcoma (ARMS) and Embryonal Rhabdomyosarcoma (ERMS) were obtained from the St. Jude Cloud Genomics Platform (Downing et al., 2012; McLeod et al., 2021). Reads were converted to gene level expression using featureCount (Rsubread package, version 2.4.3) with the Rsubread package built-in annotation (NCBI RefSeq annotation for hg38, build 38.2). Differential expression analyses were carried out between ARMS with PAX3-FOXO1 fusion biomarker (n=21) and ERMS (n=43) using DESeq2 (version 1.28.1). All analyses done in R version 4.0.0.

#### QUANTIFICATION AND STATISTICAL ANALYSIS

All analytical and statistical tests for ChIP-seq and ATAC-seq data are outlined above. The remaining tests were performed using R (3.6.1).

